# Experimentally induced drought and growing season stage modulate community-level functional traits in a temperate grassland

**DOI:** 10.1101/2023.01.12.523738

**Authors:** E. Fenollosa, P. Fernandes, A. Hector, H. King, C.S. Lawson, J. Jackson, R. Salguero-Gomez

**Author notes:** **Corresponding authors:** Erola Fenollosa, Rob Salguero-Gómez. shared senior-author (J. Jackson, R. Salguero-Gomez).

## Abstract

1. Extreme precipitation events are expected to become more intense and frequent with climate change. This climatic shift may impact the structure and dynamics of natural communities and the key ecosystem services they provide. Changes in species abundance under these extreme conditions are thought to be driven by functional traits, morpho-physiological characteristics of an organism that impact its fitness. Future environmental conditions may, therefore, favour different functional traits to those in present-day communities.
2. Here, we measure functional traits on 586 vascular plants in a temperate grassland where precipitation has been experimentally manipulated for six years. We calculated community-weighted means of five functional traits (plant height, leaf dry matter content, leaf thickness, specific leaf area, and leaf phosphorus concentration) and compared community-weighted means between three levels of precipitation: drought (−50%), irrigated (+50%), and control. Additionally, we contrasted treatments at two different timings along the growing season: mid-season and late-season.
3. We expected altered community-weighted means for traits associated with a conservative use of water that will result from increased summer stress-induced intraspecific variability in the mid-season and from community composition changes in the late-season, after the field is cut, a common management action across most European grasslands.
4. In the drought treatment, we found significantly lower community-weighted mean plant height and leaf dry matter content. However, we only observed these differences after the mid-season cut. We also observed an increase in leaf phosphorus concentration in the drought treatment before the mid-season cut. A combination of changes in community composition and intraspecific variation contributed to these differences, with community composition being more important after the cut. Species with higher height, leaf dry matter content, and lower leaf thickness showed a more pronounced abundance decline at the drought plots. We observed no changes in functional traits community-weighted means in the irrigated treatment compared to those in control and drought treatments.
5. *Synthesis*. Our results suggest how the functional trait composition of grassland communities may shift under climate change-induced drought, stressing the interacting effects with growing season stages.

## Introduction

Climate change is predicted to radically alter the structure and function of biological communities worldwide (Diaz and Cabido, 1997; IPCC, 2022). Along with increasing global temperatures, climate change will increase the intensity and frequency of extreme precipitation events (Fischer et al., 2013). Changes in the precipitation regime are likely to favour certain plants over others (MacGillivray et al., 1995; White et al., 2000; Mueller et al., 2005; Lavorel et al., 2011). Species favoured under these novel environments may have different functional traits - individual’s features that affect fitness through their influence on survival, growth, and reproduction (Díaz et al., 2016; Laughlin et al., 2020; Violle et al., 2007) - to those found under previous regimes (Lavorel et al., 2011; White et al., 2000). Changes in functional traits of individuals are expected to scale through the community level to impact ecosystem functioning (Suding et al., 2008; Woodward & Diament, 1991). Besides, because the effect of a functional trait on fitness depends on the environment, climate change is expected to alter ecosystem functioning through shifts in mean community trait values (*i*.*e*. community-weighted means) in functional traits in natural communities (Lavorel et al., 2011; McGill et al., 2006).

Functional trait-based approaches provide a promising tool for predicting community responses to climate change (Lavorel & Garnier, 2002; Quétier et al., 2007; Brodribb, 2017). However, the ability to predict changes in ecosystem functioning from environmental changes via changes in traits is considered one of the main challenges in ecology (Funk et al., 2017; Lavorel & Garnier, 2002; Suding & Goldstein, 2008). Functional traits have so far fallen short of fulfilling these ambitions (Shipley et al., 2016; Green et al., 2022). Identifying response traits, those that respond strongly to environmental gradients, is critical to predict changes in ecosystem structure and functioning in the context of climate change (Andrew et al., 2022; Lavorel & Garnier 2002; Lavorel et al., 2011; McGill et al., 2006).

Much remains unknown regarding how changes in precipitation under climate change will affect the functional traits of plant communities. Observational studies using natural precipitation gradients have shown significant correlations of functional traits along the environmental gradient (*e*.*g*., specific leaf area; Dwyer et al., 2014; Harrison et al., 2015; Wright et al., 2005). These studies, however, often struggle to attribute changes to specific environmental drivers, highlighting the need for experimental approaches (Hoover et al., 2014). However, most experimental studies to date have focused on the effect of precipitation on ecosystem functioning rather than explicitly investigating the role of functional traits in mediating community changes (Grime et al., 2000; Hoover et al., 2014; Jamieson et al., 1998; Kröel-Dulay et al., 2022). Furthermore, whether differences in community-level functional traits are driven by changes in community composition, intraspecific variation, or both, remains unknown. Failure to account for intraspecific variation is one reason why functional trait ecology has fallen short of fulfilling its predictive potential (Shipley et al., 2016; Yang et al., 2020). Indeed, functional traits vary significantly within species (Violle et al., 2012; Siefert et al., 2015; Moran et al., 2016), potentially shifting mean community trait values even if species composition remains unchanged (Pichon et al., 2022; Bricca et al., 2022). The contribution of community composition and intraspecific variation to functional traits of plant communities may also change during the growing season. For instance, early-successional communities are more sensitive to environmental changes (Grime et al., 2000; Odum, 1969). However, the interacting effects of climate gradients and vegetation developmental stage during the growing season are very complex and evidence is still scarce (Vitra et al., 2019).

To study the effect of precipitation on grassland community-level functional traits, we experimentally manipulated precipitation at RainDrop, a natural grassland near Oxford, UK. We characterised the functional trait composition in plots receiving a drought (−50% precipitation) or irrigated treatment (+50% precipitation), *vs*. control plots, which recorded the background precipitation. We calculated community-weighted means by measuring five functional traits that relate to the leaf economics spectrum and plant height, the two main axes of variation in the global spectrum of plant form and function (Díaz et al., 2016) on the most abundant species in each treatment. To measure changes in functional trait composition through the growing season, we repeated the measurements in the mid *vs*. late growing season, after a cut of the field site, a common practice across most European grasslands. We used these data to test the following hypotheses: (H1) community-weighted mean trait values will differ between precipitation treatments due an increased presence of traits associated with a more conservative use of water (Pérez-Harguindeguy et al., 2013). Specifically, we expect mean height and specific leaf area to be lower in the drought treatment with increases in leaf dry matter content and leaf thickness, and converse effects in the irrigation treatment. Leaf phosphorus concentration will decrease in the drought treatments because this trait typically correlates with specific leaf area along the leaf economics spectrum (Wright et al., 2004); (H2) community composition will differ between precipitation treatments because species with traits advantageous in drought (*e*.*g*. higher leaf thickness; Pérez-Harguindeguy et al., 2013) will increase in relative abundance in the drought treatment, with converse effects in the irrigation treatment; (H3) there will be a substantial stress-induced intraspecific variability due to species ability to adjust their physiological strategy to novel environments (Helsen et al., 2017) and contribute to community mean trait values (Hoover et al., 2014; Pichon et al., 2022), with species showing lower height and specific leaf area and higher leaf dry matter content and leaf thickness in the drought treatment, for the same reasons in H1; and (H4) community-weighted means treatment differences in the mid-growing season will be mainly influenced by increased intraspecific variability due to the annual maximum temperatures in this period (July) contributing to stress-induced variability (Helsen et al., 2017), whereas in the late-growing season community composition change may be determinant for community-weighted means due to community regrowth after the seasonal cut of the field site, as early-successional communities are more sensitive to environmental changes (Grime et al., 2000).

## Material and Methods

### Study site

The experiment was located at the Upper Seeds field site (51°46’16.8”N 117 1°19’59.1”W) in Wytham woods, Oxfordshire, UK. This is a calcareous grassland ecosystem and is managed with cuts twice per year. The first mowing takes place mid-growing season (at the end of July), and the second mowing takes place at the end of the growing season (at the end of September). To measure changes in traits through time (H4), we collected data in two different parts of the growing season; we performed initial fieldwork mid-growing season (July 2021) and again in the late-season (September 2021), just before each seasonal cut. The site has a low average soil depth (300-500 mm), generally alkaline soils (Gibson & Brown, 1991), a daily average temperature range of -5 °C to 26 °C (2016-2020), a daily total precipitation range of 0-40 mm (2016-2020). We experimentally manipulated precipitation levels using the RainDrop long-term ecological experiment which forms part of the DroughtNet global coordinated research network (https://drought-net.colostate.edu/). The experiment has been running since 2016 and consists of 25 5m^2^ plots distributed across the grassland. Each plot receives one of three precipitation treatments: drought (−50% rain), irrigated (+50% rain), and control (no manipulation). Rain shelters intercepting 50% of rain simulate drought. This rainwater is intercepted by gutters and collected in deposits situated next to each shelter. Pipes connect these deposits to sprinklers that spray the water onto an adjacent plot. This forms the irrigated treatment. This design ensures that the precipitation that each experimental treatment receives is proportional to the average natural precipitation across the site. A further set of plots undergo no precipitation manipulation and serve as ambient control plots. Additionally, to control for shelter effects, each block has one procedural control plot. These consisted of rain shelters with inverted gutters, allowing 100% of precipitation to pass through. However, previous work at this field site revealed no differences in community composition between the procedural and ambient controls (John Jackson, Personal communication). Therefore, we did not measure traits from the procedural control plots, focussing sampling effort on the precipitation treatments and ambient control plots. Each treatment is replicated across five blocks, with each block consisting of one drought plot, one irrigated plot, two ambient controls, and one procedural control. The experimental manipulations (drought and irrigation) are applied only during the main growing season (May – September). Within each block, treatments are randomly assigned with the only restriction being that the drought and irrigated plots must be next to each other for logistical reasons. To minimise edge effects, we split the 5m^2^ plot into four quarters and marked out a 1m^2^ quadrat in the centre of the study quarter from which we made all trait and abundance measurements.

### Abundance counts

To obtain abundance data to further calculate community weighted means (H1) and evaluate community composition dissimilarity between precipitation treatments (H2), we quantified species-level percentage cover for all vascular plant species using a 1m^2^ gridded quadrat (10 cm grid) in each plot. We estimated the percentage cover independently for every species in each quadrat, then transformed raw abundance data into relative abundances that sum to 100%. Because the mid-season cut removed the inflorescence from all grasses, species ID was not possible for many graminoid species during the late-season period, which may impact the observed community effects. Two graminoid species (*Brachypodium pinnatum* and *Brachypodium sylvaticum*) were identifiable to species-level because of their distinctive leaves. For these two species, we recorded percentage cover as normal. Separately, we recorded the pooled abundance of all other graminoid species. Although we identified graminoid species in the mid-season, in order to ensure a like-for-like comparison between mid-season and late-season abundance data, we combined the mid-season abundance of non-Brachypodium graminoids prior to analysis to matched the way in which we recorded the abundance of non-Brachypodium graminoids in the late-season.

### Trait measurement

To test how functional traits of grassland communities (H1) and species (H3) respond to changes in precipitation, we measured height, specific leaf area, leaf thickness, and leaf dry matter content on the most abundant species in each quadrat. For the selected species to be representative of the community, we aimed to sample species with a cumulative abundance of at least 80% within each quadrat, following Garnier et al. (2004) and Pakeman & Quested (2007). Having selected the species to be sampled in each quadrat, we randomly selected three individuals per species for trait measurement. We measured traits using standardised protocol (Pérez-Harguindeguy et al., 2013), summarised briefly in **Table 1**. We selected mature, healthy individuals where possible. For measuring leaf traits, we sampled one young but fully developed leaf per individual and measured all leaf traits on the same leaf.

**Table 1.**
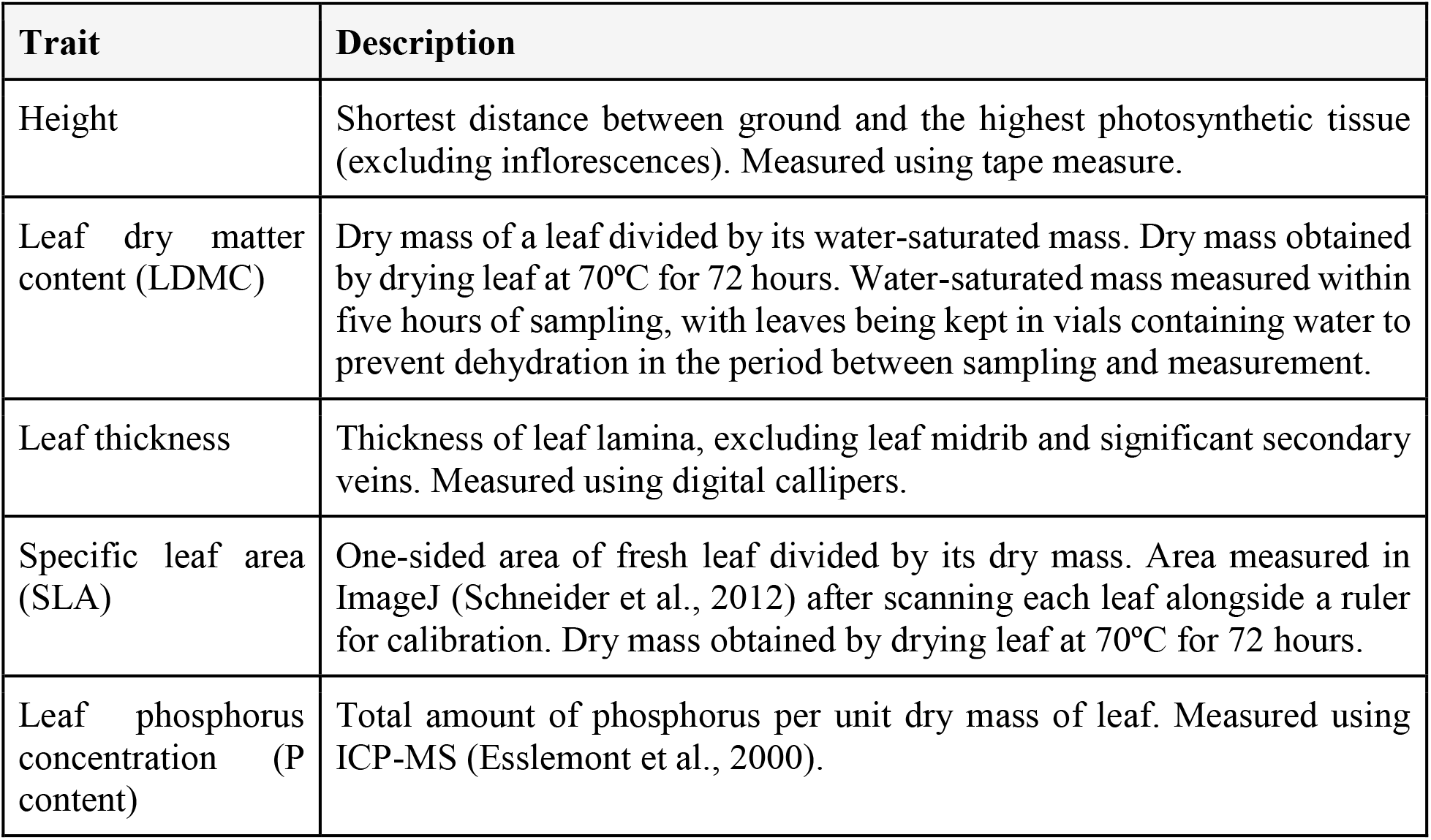
Brief descriptions of functional traits measured in this study and measurement protocol. Descriptions summarised from Pérez-Harguindeguy et al., (2013).

To test how leaf nutrient content responds to the precipitation treatments, we measured leaf phosphorus concentration using inductively coupled plasma mass spectrometry (ICP-MS). Because this technique requires 50 mg of material, we could only perform this analysis on a subset of all leaf samples. We pooled together the three replicate leaf samples per species per plot and performed the analysis on all samples where the pooled mass was at least 50 mg. After pooling, we obtained 69 samples eligible for analysis. We measured out 50 mg of each of these samples before digesting with 1 mL of concentrated nitric acid and 0.7 mL of hydrogen peroxide at 50ºC overnight. We diluted the resulting solution 25 times with MiliQ water before performing the ICP-MS (Esslemont et al., 2000).

### Community-weighted means

To compare community-level trait values between the different treatments (H1), we calculated the community-weighted mean of each trait in each quadrat. Community-weighted means are widely used to quantify shifts in community mean trait values due to environmental selection (Garnier et al., 2004). We calculated community-weighted means by multiplying the mean trait value per species in each treatment (from all collected individuals) by each species’ relative abundance in the quadrat and summing the products across all species. We rescaled the relative abundances after removing species for which no trait data was collected, following de Bello et al. (2021). We assigned trait values to each species by taking the mean across all replicates of each treatment in order to reach our target of sampling species with a cumulative abundance of 80% in almost all quadrats (**Table S1**).

Because of the mid-season cut, it was not appropriate to use a mid-season height value in the calculation of a late-season community-weighted mean, and vice versa. We therefore used different values for height for each part of the growing season. Because the other traits did not vary across the growing season (**Table S3**), we used the same trait values for both growing season stages. We used the height of *Brachypodium pinnatum* as the height of late-season graminoids as *B. pinnatum* was the only graminoid for which we measured late-season traits. For all other traits, we used the mean mid-season values across all graminoid species as the trait values of the late-season graminoids.

### Statistical analysis

We analysed data in R (R Core Team, 2021), fitting mixed-effects models using the package *lme4* (Bates et al., 2015), analysing community composition using the package *vegan* (Oksanen et al., 2020) and performing principal component analysis using the package *PCAtools* (Blighe & Lun, 2022). When interpreting the output of mixed-effects models, we opted to focus on differences based on the 95% confidence intervals rather than relying on p-values. We did this because we felt that confidence intervals provide better information about the precision of our results (Flechner & Tseng, 2011). Additionally, use of p-values has generally been discouraged with mixed-effect models (Bates et al., 2015). We considered responses to be significant if there was no overlap in the 95% confidence intervals of the treatment (drought or irrigated) and controls, highlighting cases of borderline significance.

#### Does experimentally manipulated *precipitation* alter community-weighted functional traits?

We fitted a series of models to compare community-weighted means between the precipitation treatments (H1) at the different points of the growing season (H4). To account for the blocked experimental design, we fitted hierarchical linear mixed-effect models to our data using maximum likelihood. We treated precipitation treatment and period of the growing season (mid-*vs* late-season) as fixed effects and used a hierarchical random effect structure of treatment within block. To comply with the assumptions of linear modelling, we log-transformed height, with all other trait data remaining untransformed. We fit four models that explored different combinations of fixed effects and their interactions. This involved fitting models that included only one fixed effect (treatment and growing-season period separately), both fixed effects, and both fixed effects with their interaction (**Table S2**). We compared these models to a base model which only fitted an intercept by comparing their Bayesian Information Criterion (BIC).

#### Do changes in community composition contribute towards community-weighted functional traits changes?

To assess changes in community composition between the treatments (H2) at the different growing season stages (H4), we performed non-metric multidimensional scaling (NMDS). NMDS is a form of dimension reduction that allows for differences in communities to be visualised. It is based on the rank-order of species abundances and aims to maximise the correlation between real-world distance and distance in the ordination space. We assessed the significance of any differences in community composition using analysis of similarity (ANOSIM). We used similarity percentage (SIMPER) analysis to determine which species were responsible for any dissimilarities between the treatments.

#### Does intraspecific variation contribute towards community-weighted functional traits changes?

To measure intraspecific differences within individual species (H3) in the two parts of the growing season (H4), we fit a further set of linear mixed-effects models for each species separately. Here, we fit models with only precipitation treatment as a fixed effect. As with our models for community-weighted means, we used a hierarchical random effect structure of treatment within block in all models. For every species that we analysed, we fit a separate model for each trait. Because of the mid-season cut, we analysed height separately for the mid-season and late-season stages.

## Results

Overall, we collected functional trait data on 586 individual plants across both growing season stages. These samples belonged to 22 different species in the grassland community. For most of the traits, we successfully sampled species with a cumulative abundance of 80% in 70% of quadrats (**Table S1**). The exception was leaf phosphorus concentration, where we achieved the 80% threshold in only 25% of quadrats across both growing season stages (**Table S1**).

### Community-weighted means

Precipitation treatment affected community-weighted mean trait values, but the effects were observed more strongly in the estimated values at the late-season (post-cut) than in the mid-season (pre-cut) (**Figure 1**). In the mid-season, community-weighted mean height, leaf dry matter content, leaf thickness, and leaf phosphorus concentration were higher in the drought plots compared to control, with only specific leaf area decreasing (**Figure 1**). However, only the change in leaf phosphorus concentration had non-overlapping 95% confidence intervals. In the irrigated treatment, estimates for height, leaf dry matter content, specific leaf area, and leaf phosphorus concentration were higher than in the control plots, whilst leaf thickness was lower. None of these changes had non-overlapping 95% confidence intervals.

**Figure 1.**
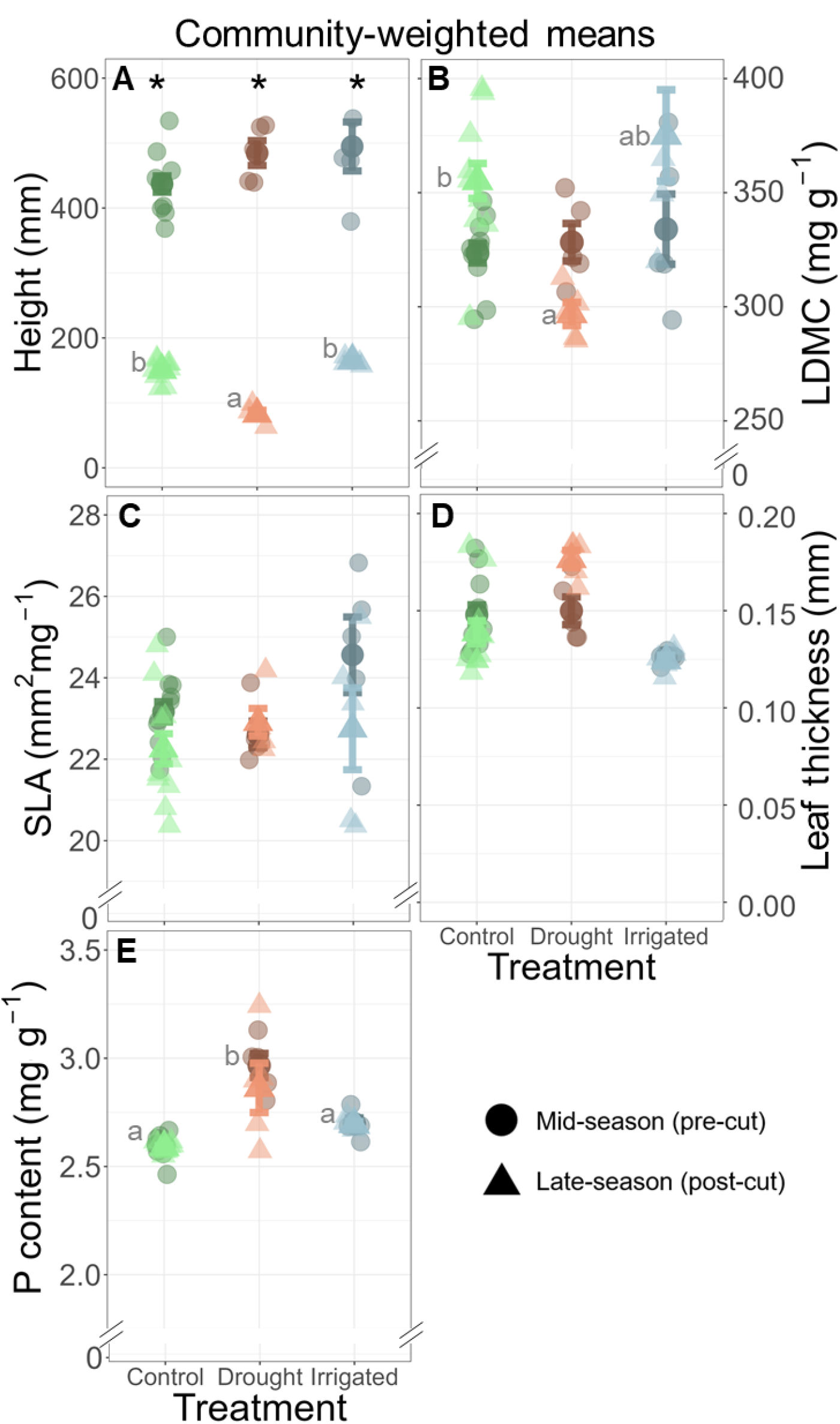
Functional traits community-weighted means shifts with treatment and growing stage. Comparison of community-weighted mean values for each functional trait (A-E) between the three precipitation treatments: control (green), drought (orange), and irrigated (blue). Each translucid small point represents the community-weighted mean of an individual plot, whilst bold points represent the mean for each period (mid-*vs*. late-season) ± SE. LDMC: leaf dry matter content; SLA: specific leaf area. Asterisks and different small letters symbolise non-overlapping 95% C.I. between stages (mid *vs*. late growing season) within each treatment and between treatments within each stage respectively. Note that in panel E, although data from different stages overlap, only significant differences between treatments were found at the mid-season stage.

In the late-season, community-weighted mean leaf thickness, specific leaf area, and leaf phosphorus concentration were higher in the drought plots compared to control, whilst height and leaf dry matter content decreased (**Figure 1**). Only the changes in height and leaf dry matter content were outside the 95% confidence intervals, with leaf thickness having very slightly overlapping intervals (control: 0.139 mm, CI 0.126 - 0.152; drought: 0.175 mm, CI 0.151 - 0.198). In the irrigated plots, leaf dry matter content and leaf thickness were higher than control plots, with height and specific leaf area decreasing and no change in leaf phosphorus concentration. None of these changes had non-overlapping 95% confidence intervals.

Only community weighted means for plant height showed non-overlapping 95% confidence intervals between growing season stages (**Figure 1A**). As expected following the cut and in the later part of the growing season, all precipitation treatments showed reduced plant height values (ca. 15cm) at the late-season (after seasonal cut) in comparison to pre-cut height values (ca. 50cm).

### Changes in community composition

One of the components of community-weighted means, community composition was different at plots with reduced precipitation in contrast with control and irrigated plots at the late-season. When plots from both growing season stages are looked at simultaneously (n = 45), there is little difference in the community composition between the treatments. This is shown by the overlapping groups in the NMDS plot (**Figure 2a**). The analysis of similarities (ANOSIM) revealed that both field work periods and treatments configure different groups. This led us to analyse the communities from each field work period (mid-vs late-season) separately. Community composition in the drought treatment cluster separately in the late-season, (n = 20, p = 0.002), but not in the mid-season (n = 25, p > 0.05) (**Figure 2b, c**).

**Figure 2.**
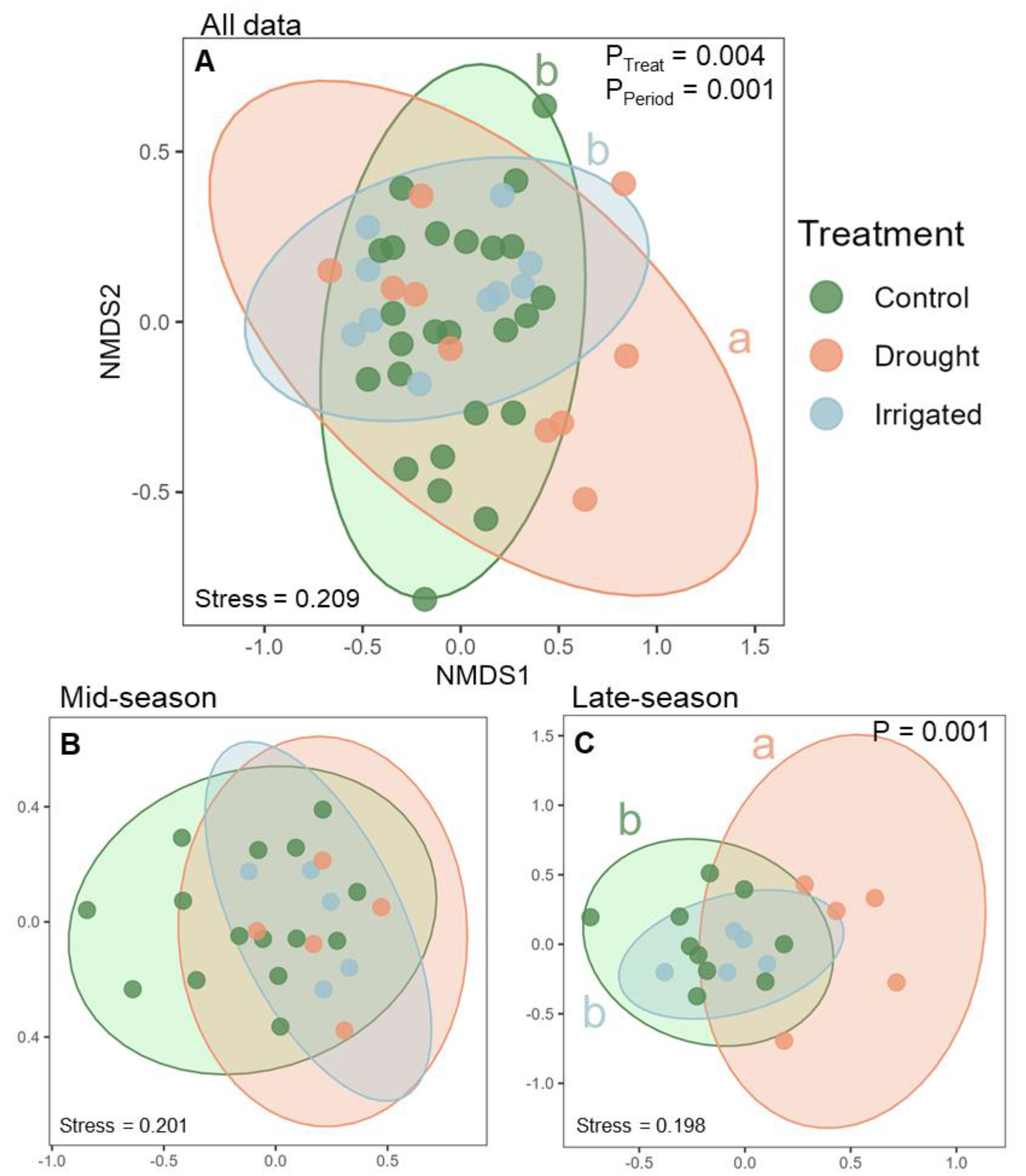
Community reassembly with precipitation treatments and growing stage. Non-metric Multi-Dimensional Scaling (NMDS) plots showing differences in community composition between the treatments. Each point represents the community composition of a single quadrat, while its location in the plot represents its position in two-dimensional ordination space. Points that are closer together are expected to have similar community composition. Stress, a measure of goodness of fit that MDS tries to minimize, is estimated as the disagreement between observed distance and ordination distance that varies between 0 (total agreement) and 1 (total disagreement), is shown at bottom left of each plot. P-values correspond to ANOSIM results for the different grouping factors: treatment and growing stage. Plots are drawn separately for (A) all community data across the summer of 2021, (B) the July 2021 communities, and (C) the September 2021 communities. Ellipses depict 95% confidence levels.

Focusing on the differences between the control and irrigated treatments with drought treatments in the late-season, similarity percentage (SIMPER) analysis revealed that three species were responsible for 70% of this difference (**Table 2**). For the comparison between control and drought treatments, these species were *Brachypodium pinnatum* (tor grass), non-Brachypodium graminoids, and the legume *Lotus corniculatus* (bird’s-foot trefoil, **Table 2**). For the comparison between irrigation and drought treatments the only difference was *Trifolium repens* (white clover), which had a higher contribution to treatment dissimilarity than *L. corniculatus*. In both comparisons, the graminoids (including *B. pinnatum*) and *T. repens* had a lower relative abundance in drought plots compared to controls whilst *L. corniculatus* had a higher relative abundance (**Table 2**). However, only the difference in graminoids abundance (including *B. pinnatum*) had non-overlapping 95% confidence intervals between control and drought treatments.

**Table 2.**
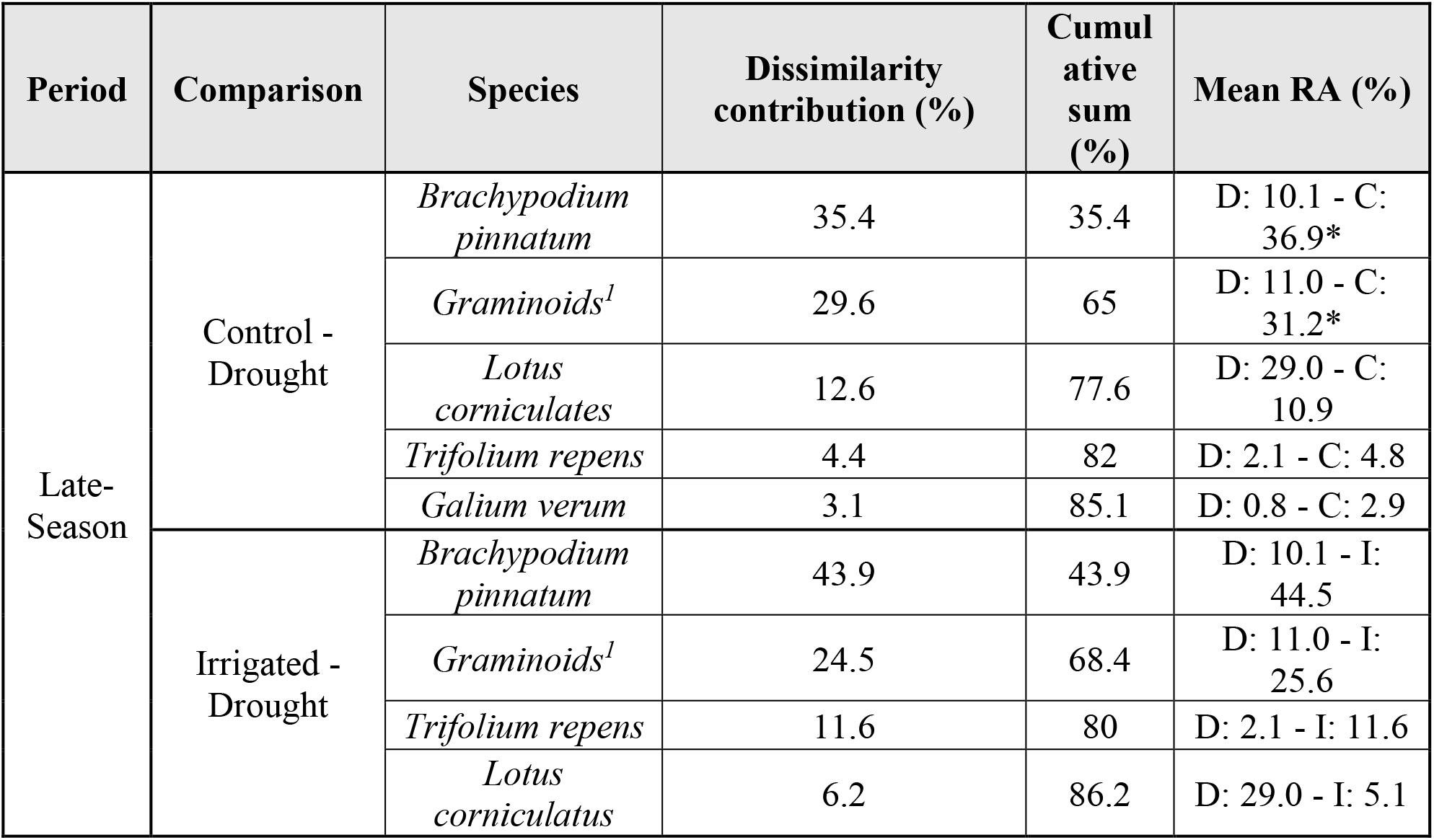
Comparison of the species that cumulatively contribute to over 85% of the dissimilarity between the differences observed in Figure 2: the communities in the drought (D) *versus* control (C) and irrigated (I) plots in the late-season. Contributions to dissimilarities were calculated using SIMPER (similarity percentage) analysis. Mean relative abundances (RA) are species absolute abundance rescaled such that the abundances of all species in a quadrat sum to 100%. The asterisk indicates non-overlapping 95% confidence intervals between relative abundances of different treatments. ^1^Excludes Brachypodium species which were assessed separately.

### Intraspecific variation

Because we sampled species depending on whether or not they were abundant in a given plot, the number of samples collected per species was not consistent. We therefore restricted analysis of intraspecific variation to the seven species with at least 30 total samples that were measured in each of the three precipitation treatments, considering the recommended replicates number to account for natural trait variation (Pérez-Harguindeguy et al., 2013). These species consisted of three graminoids (*Brachypodium pinnatum, Trisetum flavescens*, and *Arrhenatherum elatius*), three legumes (*Medicago lupulina, Trifolium repens*, and *Trifolium pratense*), and one forb (*Crepis capillaris*, **Figure 3**).

**Figure 3.**
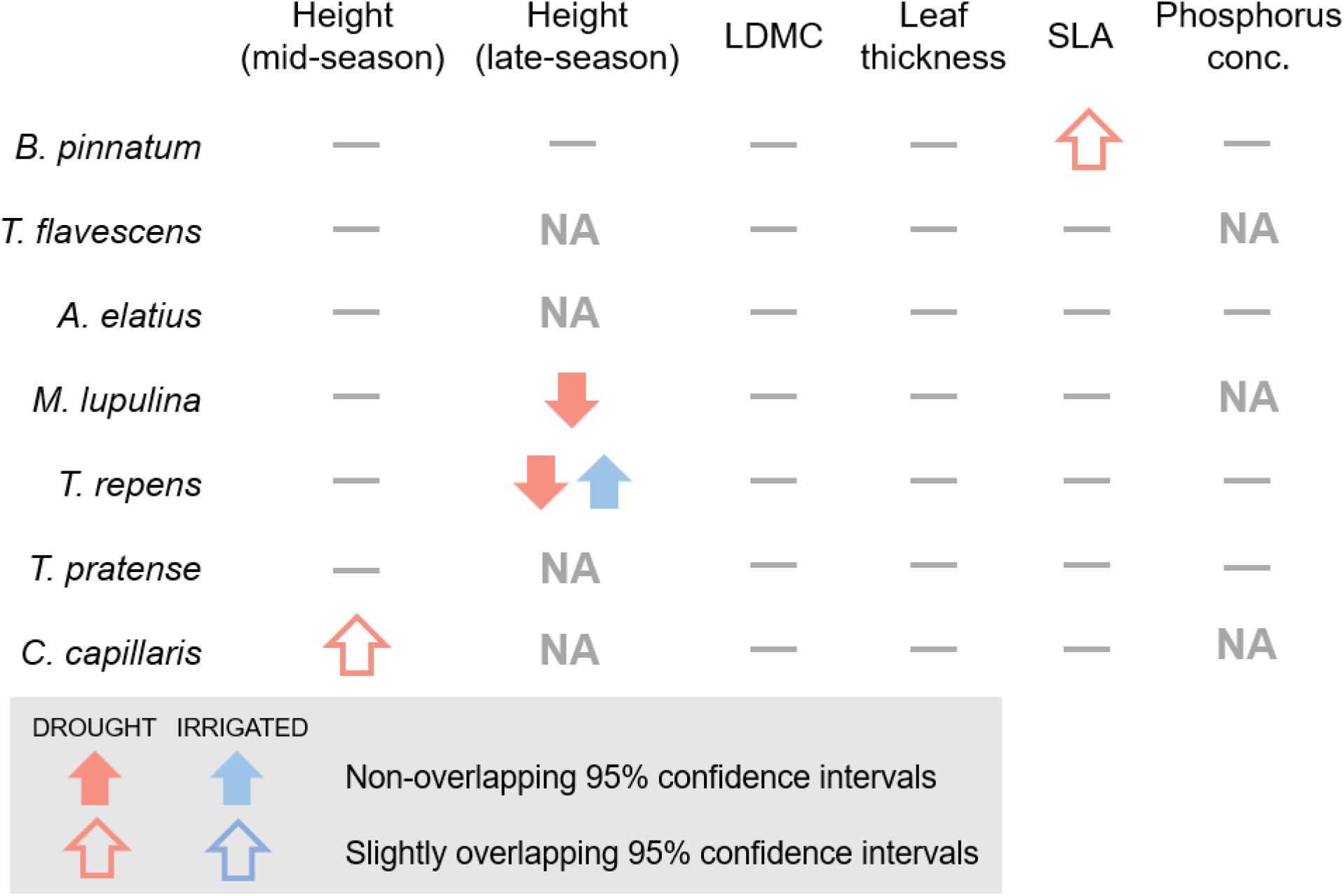
Intraspecific trait variation between precipitation treatments. Summary of intraspecific trait variation for the seven most dominant plant species in our experiment. Direction of arrows indicate change, whilst dashes represent no change in trait values. Cells marked “NA” indicate instances where no sufficient trait data were available to measure intraspecific variation. LDMC = leaf dry matter content; SLA = specific leaf area.

Of the five functional traits we measured, only height showed significant intraspecific variation between the precipitation treatments (**Figure 3**). We observed this variation only in the late-season, with two species (*M. lupulina* and *T. repens*) having a lower height in the drought treatment (*M. lupulina* control: 3.83 log(mm), CI 3.66 - 3.99; drought: 2.75 log(mm), CI 2.29 - 3.22; *T. repens* control: 4.3 log(mm), CI 4.08 - 4.51; drought: 3.74 log(mm), CI 3.41 - 4.08). *T. repens* also had a higher height in the late-season irrigated treatment (irrigated: 4.85 log(mm), CI 4.57 - 5.13). We saw a marginally significant increase in height for *C. capillaris* in the mid-season drought plots (control: 5.9, CI 5.77 - 6.03; drought: 6.14, CI 6.01 - 6.27). Other than plant height, the only other trait which varied was specific leaf area, which was marginally higher in the drought plots for *B. pinnatum* (control: 18.9 mm^2^mg^-1^, CI 17.9 - 19.7; drought: 21.8 mm^2^mg^1^, CI 19.6 - 23.8). Model output with complete means and confidence intervals for each of these seven species is summarised in **Table S3**.

When considering the four traits with most representation among species (SLA, Height, LDMC and Thickness) through principal component analysis (PCA), almost 75% of variance was explain by the two first components (**Figure 4**). The main axis of variation (PC1) was mainly positively driven by LDMC and Height and negatively by SLA, whereas Thickness was the main driver of PC2. In this bidimensional space, species with higher relative abundance differences between control and drought plots were distributed with increasing values from lower PC1 and PC2 values to higher PC1 and PC2 values. Those species that have decreased relative abundance at the drought in contrast to the control plots at the late growing season (drought sensitive species) have higher height, LDMC and may have smaller leaf thickness and SLA. For instance, this may be the case for *B. pinnatum*, that has a mean height of 610±34.5 mm, a LDMC of 433±8.30 mgg^-1^, a leaf thickness 0.11±0.002 mm and SLA of 19.2±0.32 mm^2^mg^-1^. On the opposite side of the plot, we can see more drought tolerant species, those that have increased relative abundance at the drought plots at the late growing season, such as *L. corniculatus*, which generally has high leaf thickness. This species most differentiated trait is its leaf thickness, around 0.25±0.006 mm, and a mean height of 140±17.1mm, LDMC of 245±9.43 mgg^-1^ and SLA of 21.5±1.00 mm^2^mg^-1^.

**Figure 4.**
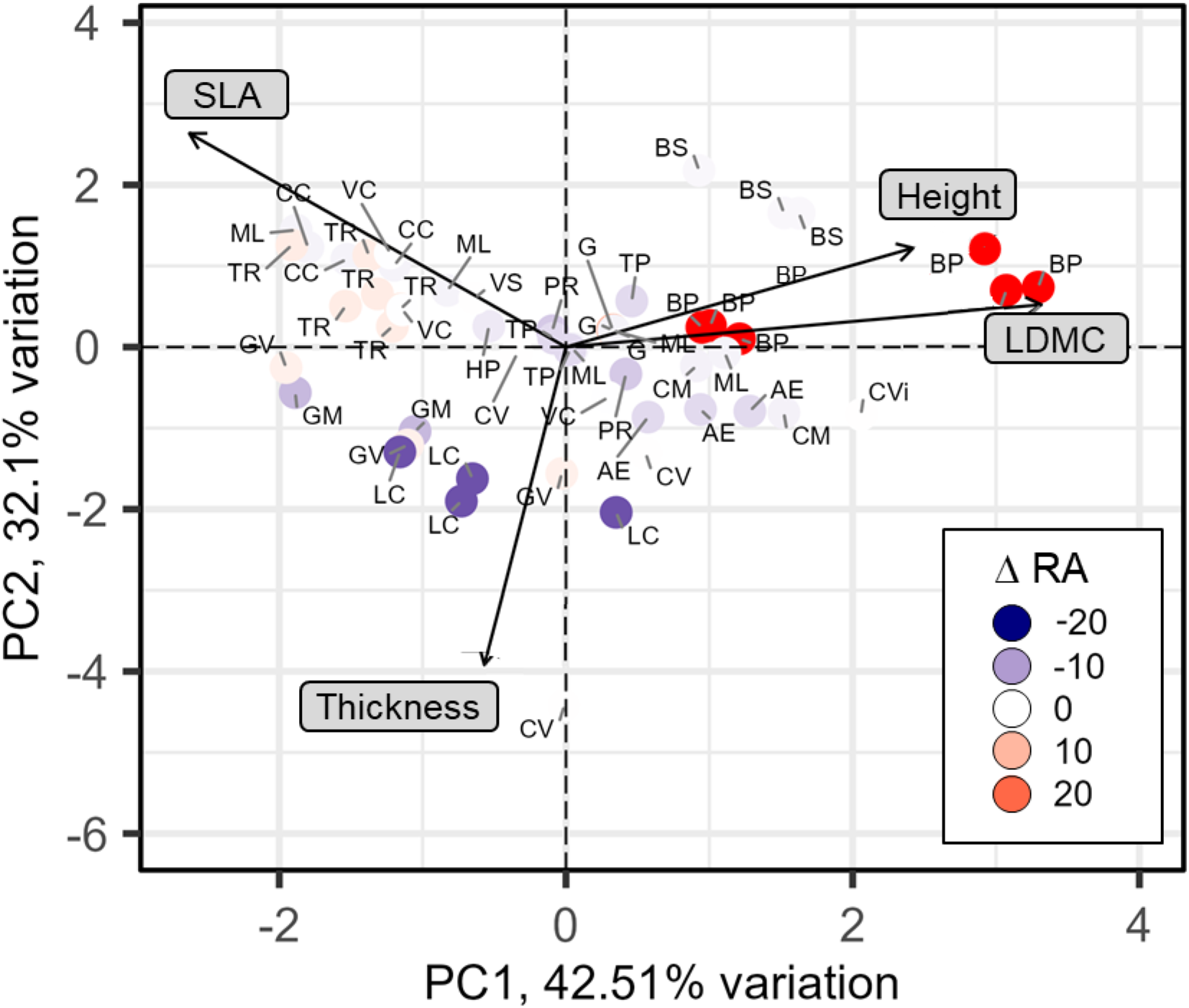
Functional traits variation plays a role on species relative abundance between precipitation treatments. Functional traits principal component analysis (PCA). Axis percentages represent the explained variance proportion from each component. The colours represent the species relative abundance difference (ΔRA) between Drought and Control treatments at the late-season (i.e., when higher differences in community composition were observed). For each species ΔRA has been calculated as mean RA in control plots – mean RA in drought plots. Species with higher ΔRA values are those that have reduced their relative abundance at drought plots. P leaf content is not included here, as only 34% of observations had all five measures, whereas considering the other four traits, 68% of the data were complete, from a total of 18 species. Each dot includes the mean value from the different plots at each treatment and growing season stage (early vs. late). Species codes are: AG: *Agrimonia eupatoria*, BP: *Brachypodium pinnatum*, BS: *B. sylvaticum*, CVi: *Clematis vitalba*, CV: *Clinopodium vulgare*, CM: *Crataegus monogyna*, CC: *Crepis capillaris*, GM: *Galium mollugo*, GV: *Galium verum*, G: *Graminoids no brachypodium*, HP: *Hypericum perforatum*, LC: *Lotus corniculatus*, ML: *Medicago lupulina*, PR: *Potentilla reptans*, TP: *Trifolium prantense*, TR: *T. repens*, VC: *Veronica chamaedrys*, VS: *Vicia sativa*.

## Discussion

In this study, we investigated how community-level functional traits respond to experimentally manipulated levels of precipitation. We found some evidence for shifting community-weighted mean functional trait values, but two traits (leaf phosphorus concentration and leaf dry matter content) varied in the opposite direction than we had hypothesised (H1). A combination of species turnover (H2) and intraspecific variation (H3) contributed to these changes. As hypothesised (H4), the relative importance of each source of variation depends on the trait in question and the different stages of the growing season (mid-vs. late-season).

### Community-level functional traits shift under drought

A 50% precipitation reduction in the temperate grassland studied communities induced shifts on three of the five studied functional traits community-weighted means. These shifts were observed according to the proposed hypothesis for plant height, but to the contrary of predictions for leaf dry matter content and phosphorous leaf content (H1). As expected, plant height community-weighted mean was significantly lower under the drought treatment, although only in the late-season. The described drought effects agree with results from observational (Fonseca et al., 2000; Moles et al., 2009) and experimental (Zuo et al., 2021) studies investigating how mean height varies along precipitation gradients in grasslands. However, no height increase was observed under the irrigated treatment. Previous work on the same field site reported high levels of evapotranspiration which may limit the effectiveness of the irrigation treatment (Jamieson et al., 1998). Plant communities at our field site may not have been water limited, meaning that individual plants would not respond to an increase in precipitation.

Contrary to our hypotheses, we observed a lower leaf dry matter content in the late-season and a higher leaf phosphorus concentration in the mid-season. Both of these hypothesised changes were originally linked to our expectation that specific leaf area would decrease in the drought treatment, as reported in observational studies (Dwyer et al., 2014; Harrison et al., 2015; Wright et al., 2005). A lower specific leaf area could have increased leaf dry matter content because of the geometric relationship between the two traits through the following equation: LDMC=1/(SLA×Leaf thickness) (Vile et al., 2005). In our results, the marginal evidence for an increased leaf thickness was countered by a lower leaf dry matter content. These joint changes cancel each other out, with the net effect being no change in specific leaf area. Likewise, specific leaf area has been shown to correlate with leaf nutrient content in the leaf economics spectrum (Wright et al., 2004). Because we did not observe the expected specific leaf area response, other processes may have led to the measured changes in leaf dry matter content and phosphorus concentration. LDMC is a measure of investment of the plant species in defence and structural components and therefore is strongly related to plant productivity (Pérez-Harguindeguy et al., 2013). A possible explanation of the observed LDMC reduction under the drought treatment could be a consequence of a delayed leaf development in the drought treatment. Indeed, LDMC is strongly related to seasonal and developmental effects, with younger leaves having lower LDMC values (Palacio et al., 2008).

In contrast to our hypotheses, we found increases in leaf phosphorus concentration under drought conditions in the mid-season. Our result matches the finding of Wright et al. (2001), who reported higher leaf phosphorus concentrations in species growing in dry sites compared to wet sites in an observational study. They explained this result in terms of a greater investment in photosynthetic enzymes, leading to a higher nitrogen concentration (a trait that scaled with phosphorus concentration). This would allow plants to achieve a high photosynthetic rate whilst maintaining low stomatal conductance, limiting water loss. Another reason we might expect to find an increased leaf phosphorus concentration in the drought treatment is that the soil nutrient content may be higher. Jamieson et al. (1998) used a similar experimental design on the same field site as the present study and measured higher levels of nitrogen mineralisation under drought conditions. They suggested that this was linked to higher inputs of leaf litter resulting from higher rates of senescence. Although we did not measure soil nutrient content, we did observe high levels of dead plant matter in the drought plots. Additionally, Sternberg et al., (1999) used the same field experiment as Jamieson et al. (1998) and showed that leaf litter was higher in the drought plots. If this effect results in higher soil phosphorus content, and P assimilation is not limited by other factors, we would expect plants growing in these plots to have a higher leaf phosphorus concentration (Wright et al., 2001). Our result of an increased leaf phosphorus concentration in the mid-season should, however, be treated with caution. Because we could only measure phosphorus concentration on the leaves with the highest mass, we could not collect data for many of the species. This trait did not meet our sampling objective of species with a cumulative abundance of 80% in any of the 20 mid-season quadrats (**Table S1**). Therefore, our values may not be representative of the communities found in each quadrat (Garnier et al., 2004; Pakeman & Quested, 2007).

The difference in community-level traits between the mid- and late-season stages may be linked to the mid-season cut of the field site. Previous studies have found that early-successional communities are more sensitive to environmental changes (Grime et al., 2000; Odum, 1969). This effect could explain our finding of more changes in community-weighted mean trait values after the cut than before the cut. Furthermore, stronger effects of drought on community-weighted functional traits were observed after the growing peak (late-season) in two permanent grassland experimental sites at the Swiss Jura Mountains, which coincide with the longer and warmer summer days (Vitra et al., 2019). In our study, increased temperature and lower humidity may contribute to enhanced drought effects at the late-season. As highlighted by Vitra et al., (2019), the interacting effects of the timing of drought and the development stage of the vegetation during the growing season are very complex and evidence is still scarce.

### Drought induced community reassembly contributes to community-weighted functional traits shifts at the late-season

As hypothesised (H4), we found differences in community composition between the precipitation treatments in the late-season, that contribute to community-weighted functional traits shifts in the grassland community. The absence of this effect in the mid-season may be related to the increased environmental sensitivity of early-successional communities (Grime et al., 2000; Odum, 1969). Short-term (1-3 years) manipulative precipitation experiments report absent or small effects on community reassembly (Batbaatar et al., 2021; Vitra et al., 2019). In the short term (1-2 years), Vitra et al. (2019) reported that the observed changes in community-weighted functional traits in response to drought were mainly related to changes in plant traits rather than changes in species abundance (Vitra et al., 2019), instead of species turnover and community composition change, which would occur over longer drought perturbations (Smith et al. 2009). After six years of manipulated precipitation, we have observed community composition changes with an important decrease of graminoid abundance under drought. The lower abundance of grasses in drought plots agrees with other experimental studies in calcareous grasslands (Morecroft et al., 2004; Sternberg et al., 1999). Interestingly, similar effects were observed under a comparative between manipulated and observational precipitation gradients with both species turnover and intraspecific variations contributed to community-weighted functional traits responses of grass community traits to precipitation changes (Zuo et al., 2021).

### Intra and interspecific functional traits variability for drought resistance

Contrary to our hypotheses (H3, H4) and studies that have stressed the importance of intraspecific trait variation (Pichon et al., 2022; Shipley et al., 2016; Violle et al., 2012), we found limited evidence of intraspecific variation in our data. Our results found that only one of the five traits we studied (plant height) varied significantly between the precipitation treatments and in more than one species. It is possible that we did not have sufficient sample sizes to detect intraspecific variation in this study. Our sampling was primarily aimed at sampling a range of different species to calculate community-level trait values. This meant that our sampling was spread out across many species, limiting our ability to detect intraspecific changes. Non-significant intraspecific differences in trait values may still contribute to significant differences seen at the community level. This is likely the source of the mid-season increases in community-level leaf phosphorus concentration. For abundant species such as the graminoid *B. pinnatum* (tor grass) and the legume *T. repens* (white clover), there was a non-significant increase in leaf phosphorus concentration in the drought treatment. Despite this increase being non-significant when analysed at the species level, these differences may combine to form a significant difference at the community level. Therefore, intraspecific variation may be more important in determining community-level trait values than our results suggest.

Rather than intraspecific variation, interspecific differences (*i*.*e*. differences between species) were determinant for community-weighted functional traits shifts, as certain trait syndromes contribute to species drought resistance. From the perspective of functional traits, the lower abundance of grasses in drought plots is to be expected as grasses are taller plants with thinner leaves. Here, the graminoid *B. pinnatum* (tor grass) was the tallest species in the late-season and had the second thinnest leaves of all species. Those traits, together with a high LDMC may be disadvantageous in drought, explaining the grasses absence at the drought treatment. Although we did not measure root traits, the low root depth in many grass species may also contribute to their lower abundance in drought plots (Morecroft et al., 2004; Sternberg et al., 1999). The legume *Lotus corniculatus* (bird’s-foot trefoil) was notably more abundant than the grasses in the late-season drought plots. The traits of *L. corniculatus* are generally on the opposite end of the spectrum to grasses. In other words, *L. corniculatus* tends to be a short plant, with the second thickest leaves of all species measured. The relative success of *L. corniculatus* in the drought plots suggests that it has traits that are more suited to growing at low precipitation levels.

### Conclusions

Our results provide insights into how grassland communities will respond to climate change. Overall, we found evidence that short, thick-leaved plants may be favoured under extreme drought conditions, whilst grasses may become less abundant. We observed some intraspecific trait plasticity in response to drought, but the most dramatic effects were the changes in community composition. Although we did not observe changes in community structure in the mid-season, such changes are generally expected to take place over longer timescales than changes in plant morphology (Suding et al., 2008). In this context, it is perhaps not surprising that we have not yet seen changes in year-round community structure given that the experiment is still in its sixth year. The temporal variation of our community-weighted mean trait values suggests that any effect of traits on ecosystem functioning would not be consistent across time. Any trait-based attempt to predict ecosystem functioning must account for such temporal variation in community-level trait values. This may prove to be an important step towards the “Holy Grail” of predicting ecosystem functioning from changes in traits.

The trait changes that we observed may have key implications for ecosystem functioning. For example, increasing leaf thickness has been linked to a lower litter decomposition rate and lower palatability to consumers, affecting nutrient cycling and trophic interactions (Díaz et al., 2004). Lower plant height decreases carbon storage, whilst a high leaf phosphorus concentration is thought to provide a higher quality of food to consumers (Díaz et al., 2004; Moles et al., 2009). Some of these changes may cancel each other out. For example, we observed some evidence for a higher leaf thickness and higher leaf phosphorus concentration in the drought treatment (albeit at different parts of the growing season). These traits are predicted to influence processes such as litter decomposition in opposite ways (Díaz et al., 2004). In this study, we focussed on identifying response traits without simultaneously measuring ecosystem functioning. An important next step, therefore, is to verify whether the community-level trait changes that we have outlined have the expected effects on ecosystem functioning. This would strengthen predictions about how climate change-induced extreme precipitation events will impact ecosystem functioning.

## Acknowledgments

We are grateful to the technical team at Wytham Woods led by N Fisher for their valuable management of the experimental sites at Wytham Woods. Special thank C Adelmant for her for support in botanical surveys and field logistic support. EF was supported by a Margarita Salas postdoctoral Fellowship from the Ministry of Universities in Spain hosted by RSG. The analyses were supported by a Kirby Fund grant to PF. The work was supported by the Ecological Continuity Trust to AH, and a NERC Independent Research Fellowship (NE/M018458/1) to RSG.

## Authors contribution

Study design and data collection was performed by PF, JJ, RS-G, CSL, AH and HK. PF analysed the data and prepared an academic report under the supervision of RS-G and JJ. EF configured the first manuscript draft including additional data analysis. Initial manuscript feedback was provided by RS-G and JJ. Further manuscript feedback was provided by all authors, who approved the manuscript for publication.

## Supplementary material

**Table S1.**
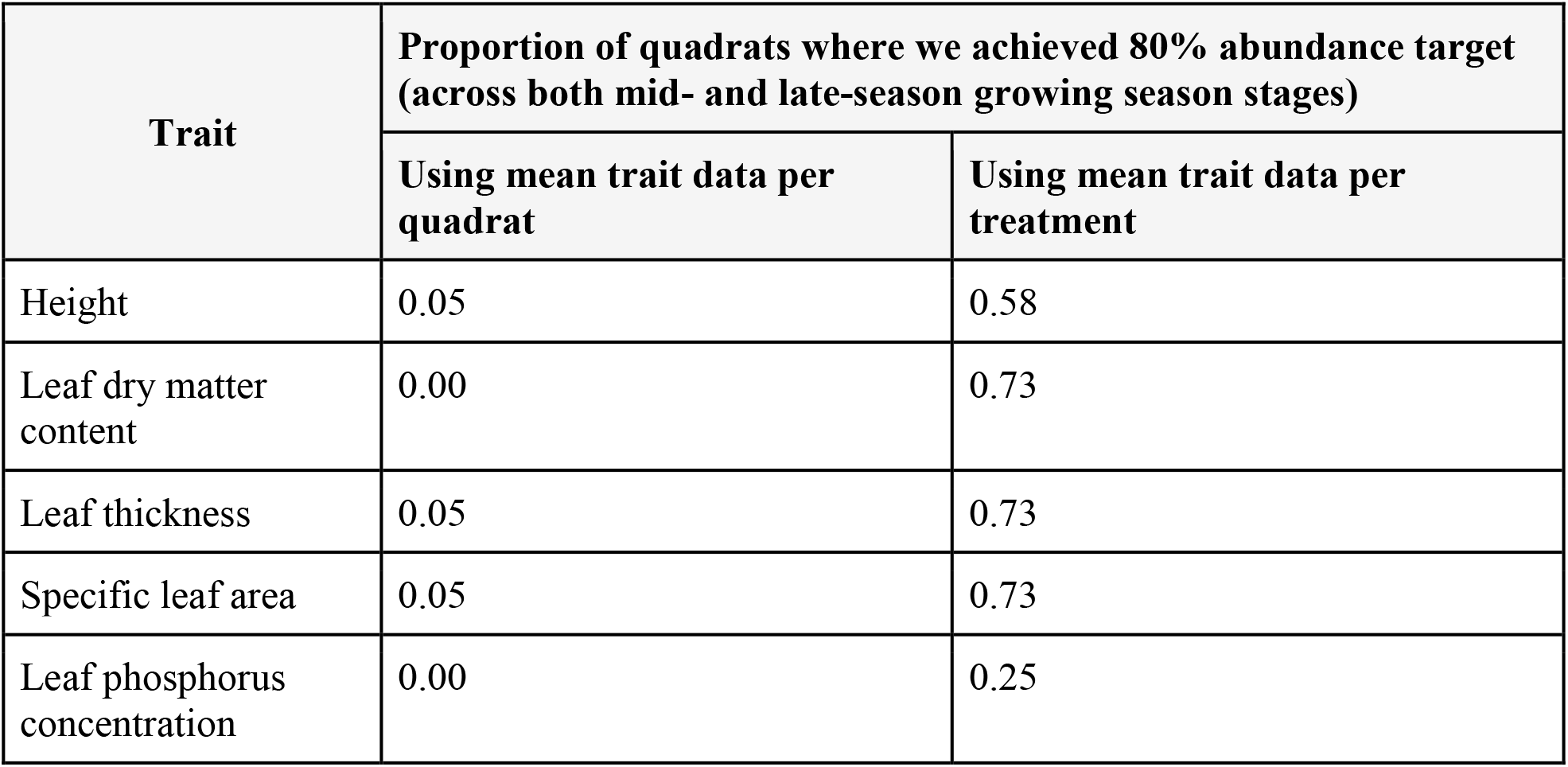
Cumulative abundance of sampled species. Proportion of the 40 quadrats (20 mid- and 20 late-season) where we achieved the 80% cumulative abundance target when selecting species for trait measurement. We compared this proportion between two different methods of calculating community-weighted trait means: 1) using the mean trait data as calculated within each quadrat and 2) using mean trait data as calculated across all replicates of each treatment. Using the second method substantially improved the proportion of quadrats that achieved the 80% cumulative abundance target. We therefore presented the results using this method in this paper. The proportion of quadrats achieving the target is not equal across all traits because there were some species for which it was not possible to measure certain traits. This mainly occurred for two reasons. Firstly, leaf phosphorus concentration could not be measured on species with light leaves because 50mg of plant material was needed for ICP-MS. Secondly, the mid-season cut meant that it was not appropriate to use the pre-cut height of a species in the post-cut calculation of a community-weighted mean, and *vice versa*. As a result, species that were sampled in only one growing season stage are missing height data for the other period.

| Trait | Proportion of quadrats where we achieved 80% abundance target (across both mid- and late-season growing season stages) |  |
| --- | --- | --- |
|  | Using mean trait data per quadrat | Using mean trait data per treatment |
| Height | 0.05 | 0.58 |
| Leaf dry matter content | 0.00 | 0.73 |
| Leaf thickness | 0.05 | 0.73 |
| Specific leaf area | 0.05 | 0.73 |
| Leaf phosphorus concentration | 0.00 | 0.25 |

**Table S2.**
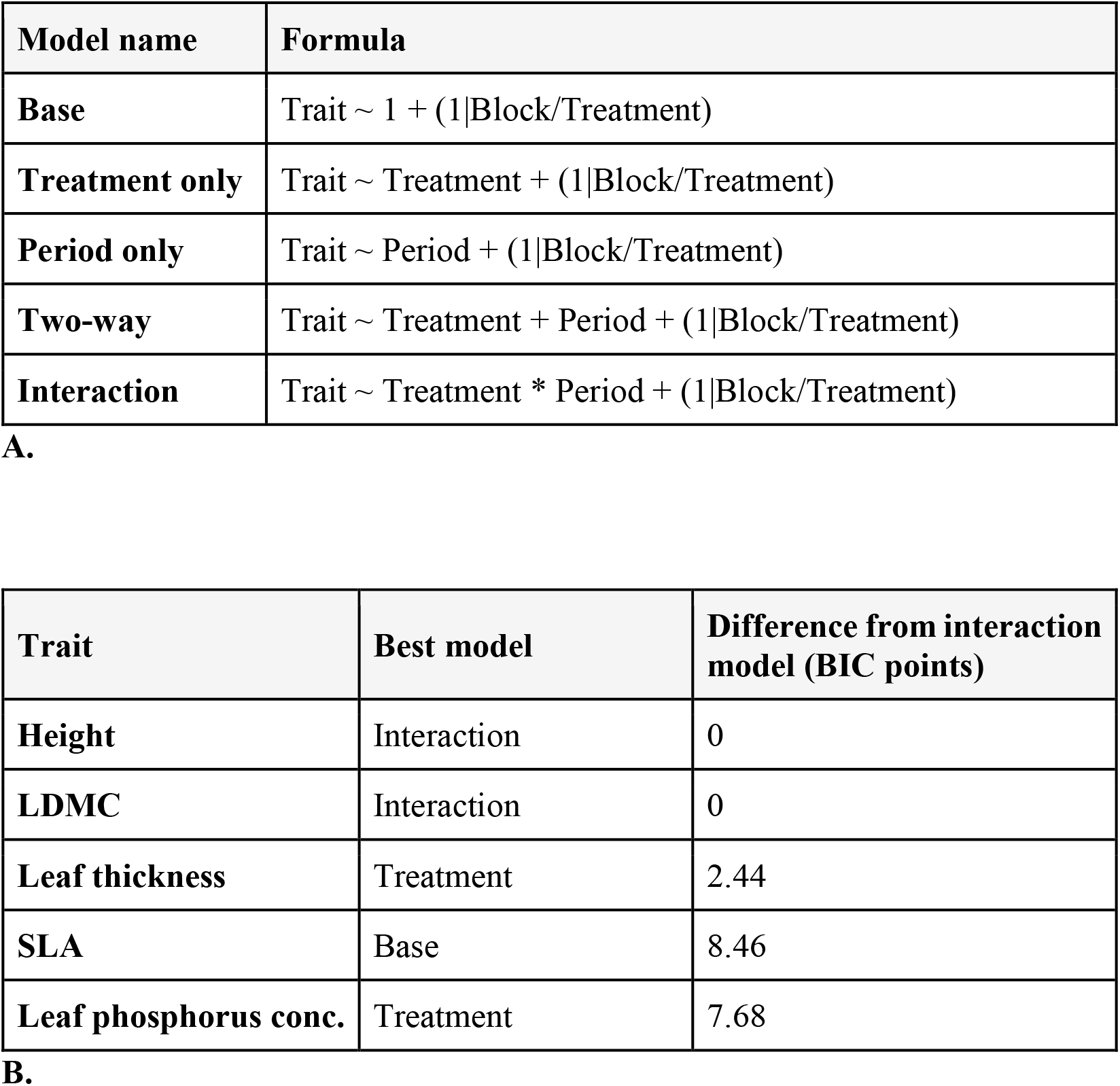
Model comparison. For fitting linear mixed-effect models’ differences to our community-weighted mean data, we explored different combinations of fixed effects. We compared models on their Bayesian Information Criterion (BIC) to inform which model we would use. Because of our experimental design, we used the same random effect structure of treatment within block for all models. Section A summarises the models that we compared. “Period” refers to the point of the growing season (mid-vs late-season). Section B summarises the best model (based on BIC) for each trait, together with the difference in BIC points between the best model and the interaction model. For height and LDMC, the model with the lowest BIC was the interaction model, whilst the treatment-only model was selected for leaf thickness and phosphorus concentration. For SLA, the base model had the lowest BIC. However, because the interaction model had a similar BIC to the best model for all traits (within 10 BIC points), it was used for all subsequent analyses.

**Table S3.**
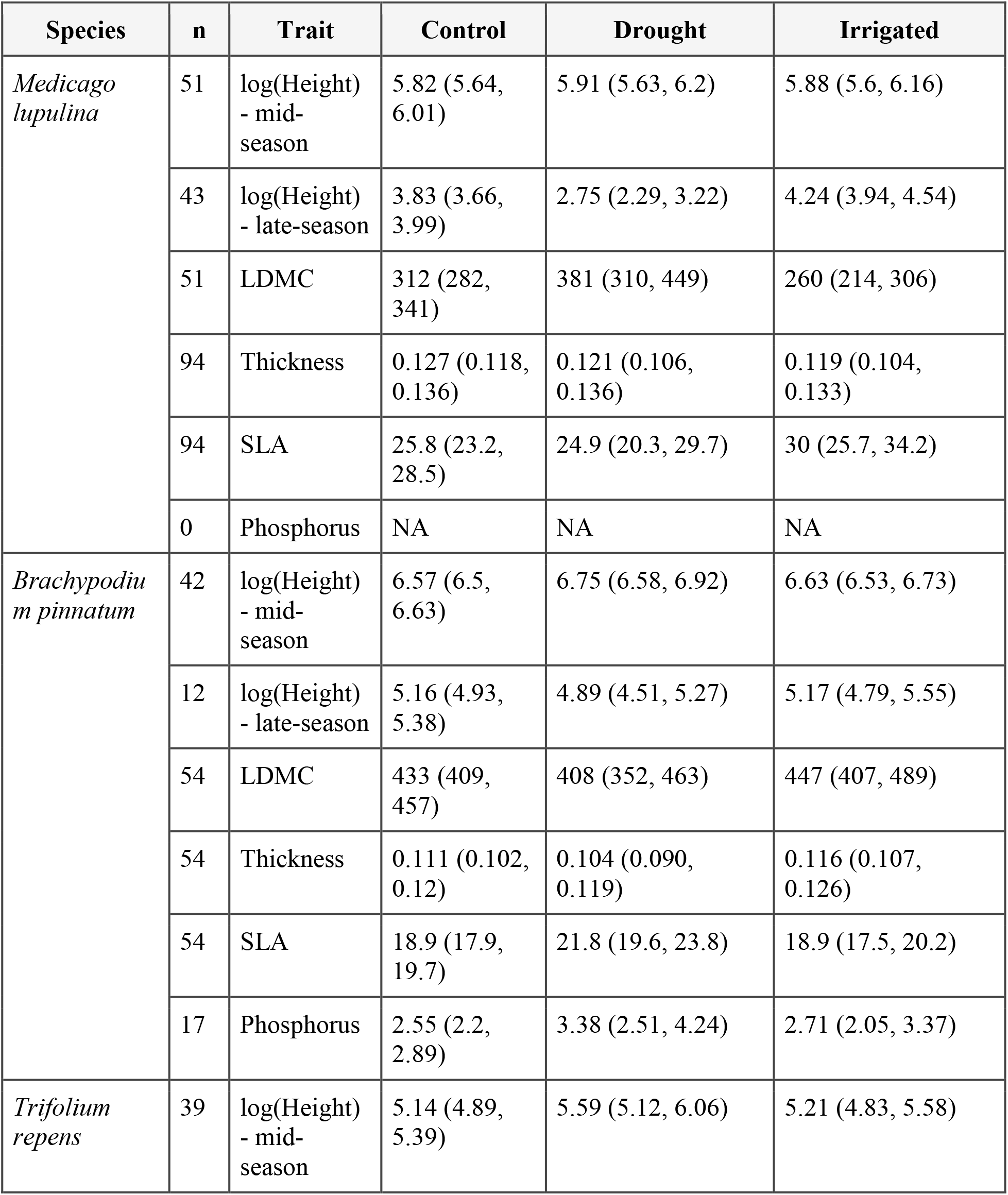
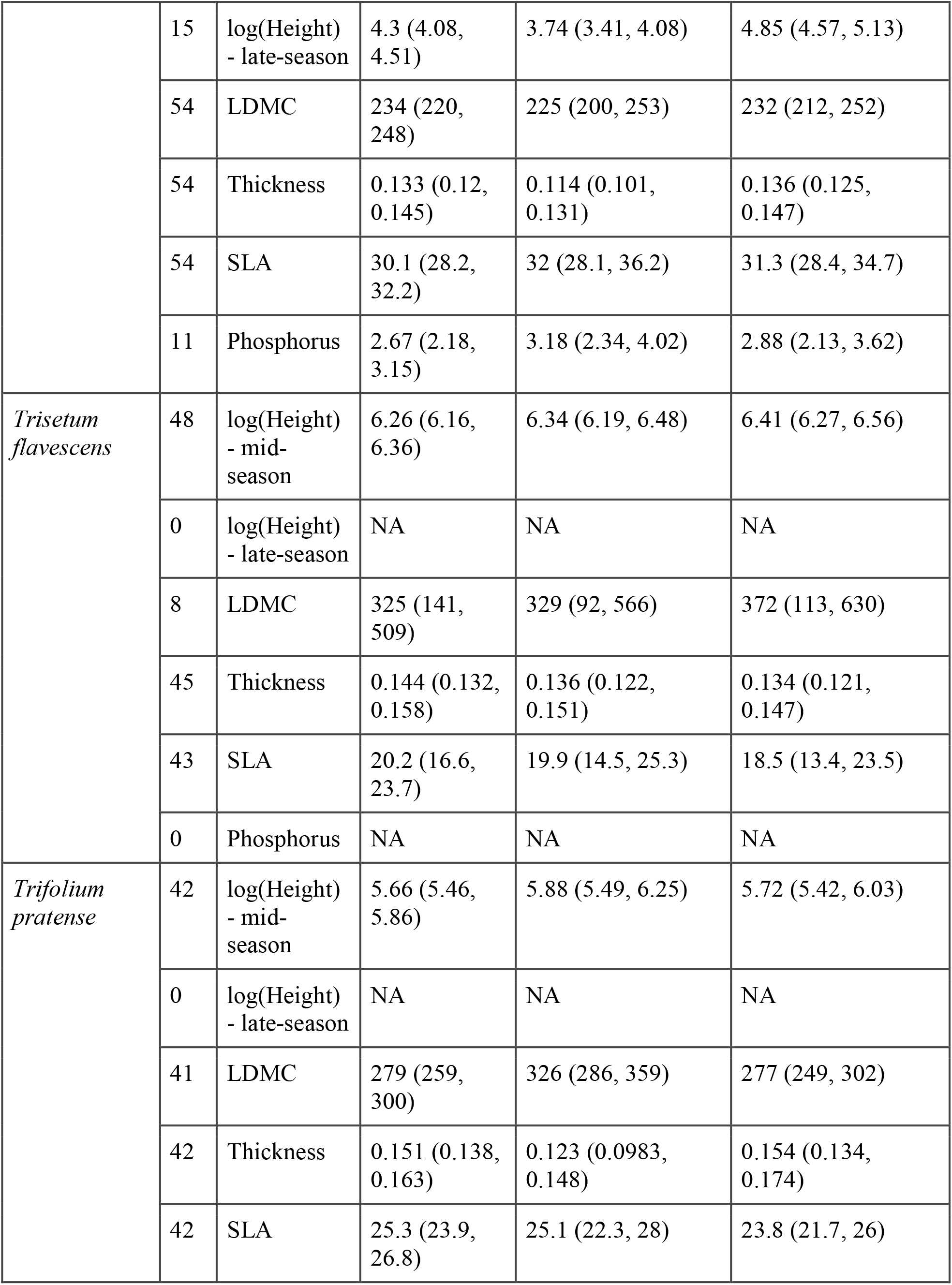
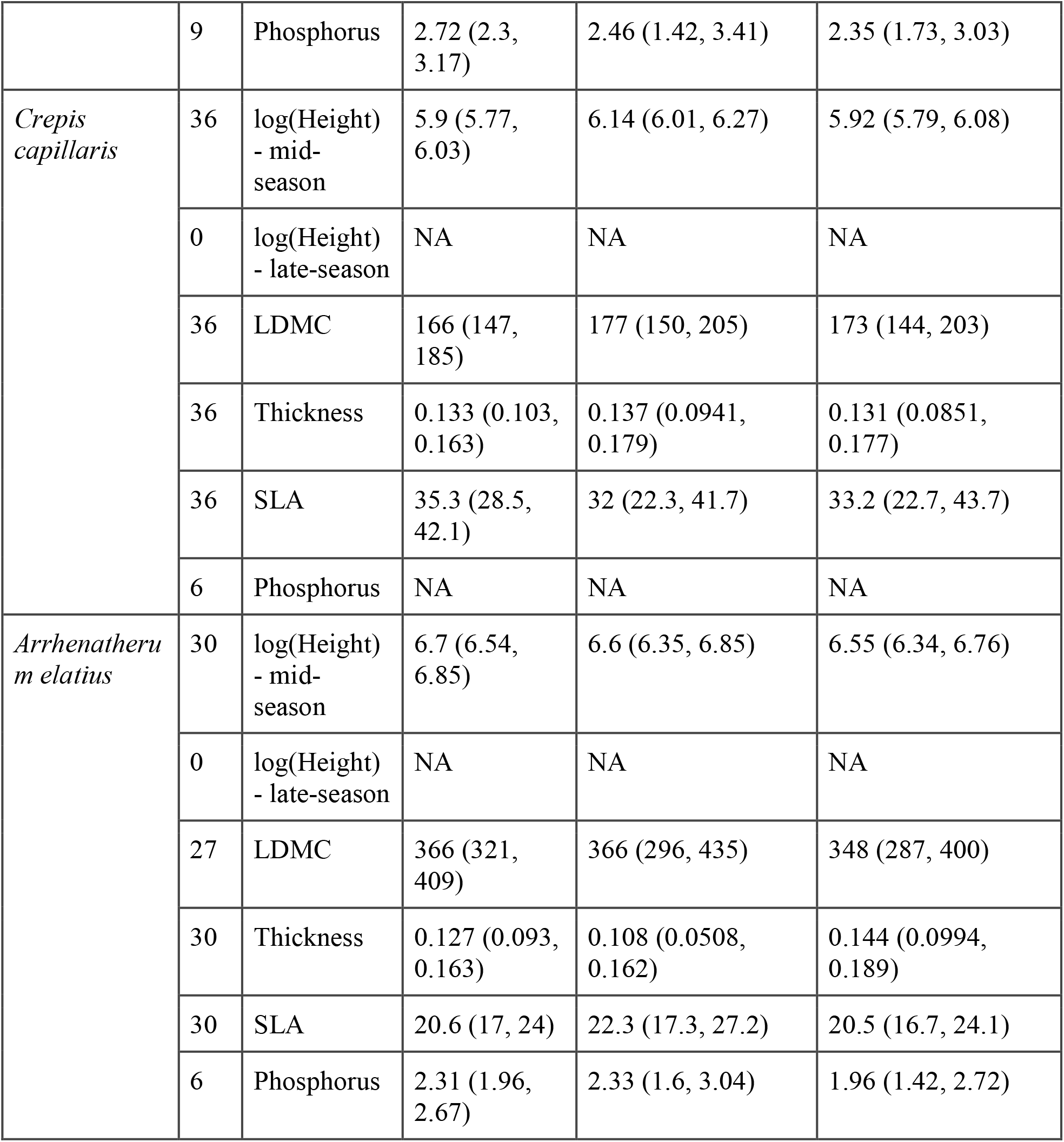
Intraspecific variation summary. Output of linear mixed-effect models investigating intraspecific variation of the seven sampled species that have at least 30 replicates. The table shows estimates for coefficients in each treatment together with 95% confidence intervals.

