## Supplementary material for "Experimentally induced drought and growing season stage modulate community-level functional traits in a temperate grassland"

**Table S2.** Model comparison. For fitting linear mixed-effect models' differences to our community-weighted mean data, we explored different combinations of fixed effects. We compared models on their Bayesian Information Criterion (BIC) to inform which model we would use. Because of our experimental design, we used the same random effect structure of treatment within block for all models. Section A summarises the models that we compared. "Period" refers to the point of the growing season (mid- vs late-season). Section B summarises the best model (based on BIC) for each trait, together with the difference in BIC points between the best model and the interaction model. For height and LDMC, the model with the lowest BIC was the interaction model, whilst the treatment-only model was selected for leaf thickness and phosphorus concentration. For SLA, the base model had the lowest BIC. However, because the interaction model had a similar BIC to the best model for all traits (within 10 BIC points), it was used for all subsequent analyses.

<table>
<tr><th>Model name</th><th>Formula</th></tr>
<tr><td>Base</td><td>Trait ~ 1 + (1|Block/Treatment)</td></tr>
<tr><td>Treatment only</td><td>Trait ~ Treatment + (1|Block/Treatment)</td></tr>
<tr><td>Period only</td><td>Trait ~ Period + (1|Block/Treatment)</td></tr>
<tr><td>Two-way</td><td>Trait ~ Treatment + Period + (1|Block/Treatment)</td></tr>
<tr><td>Interaction</td><td>Trait ~ Treatment * Period + (1|Block/Treatment)</td></tr>
</table>

**A.**

| Trait | Best model | Difference from interaction model (BIC points) |
|---|---|---|
| **Height** | Interaction | 0 |
| **LDMC** | Interaction | 0 |
| **Leaf thickness** | Treatment | 2.44 |
| **SLA** | Base | 8.46 |
| **Leaf phosphorus conc.** | Treatment | 7.68 |

**B.**

**Table S3.** Intraspecific variation summary. Output of linear mixed-effect models investigating intraspecific variation of the seven sampled species that have at least 30 replicates. The table shows estimates for coefficients in each treatment together with 95% confidence intervals.

| Species | n | Trait | Control | Drought | Irrigated |
|---|---|---|---|---|---|
| *Medicago lupulina* | 51 | log(Height) - mid-season | 5.82 (5.64, 6.01) | 5.91 (5.63, 6.2) | 5.88 (5.6, 6.16) |
| | 43 | log(Height) - late-season | 3.83 (3.66, 3.99) | 2.75 (2.29, 3.22) | 4.24 (3.94, 4.54) |
| | 51 | LDMC | 312 (282, 341) | 381 (310, 449) | 260 (214, 306) |
| | 94 | Thickness | 0.127 (0.118, 0.136) | 0.121 (0.106, 0.136) | 0.119 (0.104, 0.133) |
| | 94 | SLA | 25.8 (23.2, 28.5) | 24.9 (20.3, 29.7) | 30 (25.7, 34.2) |
| | 0 | Phosphorus | NA | NA | NA |
| *Brachypodium pinnatum* | 42 | log(Height) - mid-season | 6.57 (6.5, 6.63) | 6.75 (6.58, 6.92) | 6.63 (6.53, 6.73) |
| | 12 | log(Height) - late-season | 5.16 (4.93, 5.38) | 4.89 (4.51, 5.27) | 5.17 (4.79, 5.55) |
| | 54 | LDMC | 433 (409, 457) | 408 (352, 463) | 447 (407, 489) |
| | 54 | Thickness | 0.111 (0.102, 0.12) | 0.104 (0.090, 0.119) | 0.116 (0.107, 0.126) |
| | 54 | SLA | 18.9 (17.9, 19.7) | 21.8 (19.6, 23.8) | 18.9 (17.5, 20.2) |
| | 17 | Phosphorus | 2.55 (2.2, 2.89) | 3.38 (2.51, 4.24) | 2.71 (2.05, 3.37) |
| *Trifolium repens* | 39 | log(Height) - mid-season | 5.14 (4.89, 5.39) | 5.59 (5.12, 6.06) | 5.21 (4.83, 5.58) |

| | | | | | |
|---|---|---|---|---|---|
| | 15 | log(Height) - late-season | 4.3 (4.08, 4.51) | 3.74 (3.41, 4.08) | 4.85 (4.57, 5.13) |
| | 54 | LDMC | 234 (220, 248) | 225 (200, 253) | 232 (212, 252) |
| | 54 | Thickness | 0.133 (0.12, 0.145) | 0.114 (0.101, 0.131) | 0.136 (0.125, 0.147) |
| | 54 | SLA | 30.1 (28.2, 32.2) | 32 (28.1, 36.2) | 31.3 (28.4, 34.7) |
| | 11 | Phosphorus | 2.67 (2.18, 3.15) | 3.18 (2.34, 4.02) | 2.88 (2.13, 3.62) |
| *Trisetum flavescens* | 48 | log(Height) - mid-season | 6.26 (6.16, 6.36) | 6.34 (6.19, 6.48) | 6.41 (6.27, 6.56) |
| | 0 | log(Height) - late-season | NA | NA | NA |
| | 8 | LDMC | 325 (141, 509) | 329 (92, 566) | 372 (113, 630) |
| | 45 | Thickness | 0.144 (0.132, 0.158) | 0.136 (0.122, 0.151) | 0.134 (0.121, 0.147) |
| | 43 | SLA | 20.2 (16.6, 23.7) | 19.9 (14.5, 25.3) | 18.5 (13.4, 23.5) |
| | 0 | Phosphorus | NA | NA | NA |
| *Trifolium pratense* | 42 | log(Height) - mid-season | 5.66 (5.46, 5.86) | 5.88 (5.49, 6.25) | 5.72 (5.42, 6.03) |
| | 0 | log(Height) - late-season | NA | NA | NA |
| | 41 | LDMC | 279 (259, 300) | 326 (286, 359) | 277 (249, 302) |
| | 42 | Thickness | 0.151 (0.138, 0.163) | 0.123 (0.0983, 0.148) | 0.154 (0.134, 0.174) |
| | 42 | SLA | 25.3 (23.9, 26.8) | 25.1 (22.3, 28) | 23.8 (21.7, 26) |

| | | | | | |
|---|---|---|---|---|---|
| | 9 | Phosphorus | 2.72 (2.3, 3.17) | 2.46 (1.42, 3.41) | 2.35 (1.73, 3.03) |
| *Crepis capillaris* | 36 | log(Height) - mid-season | 5.9 (5.77, 6.03) | 6.14 (6.01, 6.27) | 5.92 (5.79, 6.08) |
| | 0 | log(Height) - late-season | NA | NA | NA |
| | 36 | LDMC | 166 (147, 185) | 177 (150, 205) | 173 (144, 203) |
| | 36 | Thickness | 0.133 (0.103, 0.163) | 0.137 (0.0941, 0.179) | 0.131 (0.0851, 0.177) |
| | 36 | SLA | 35.3 (28.5, 42.1) | 32 (22.3, 41.7) | 33.2 (22.7, 43.7) |
| | 6 | Phosphorus | NA | NA | NA |
| *Arrhenatherum elatius* | 30 | log(Height) - mid-season | 6.7 (6.54, 6.85) | 6.6 (6.35, 6.85) | 6.55 (6.34, 6.76) |
| | 0 | log(Height) - late-season | NA | NA | NA |
| | 27 | LDMC | 366 (321, 409) | 366 (296, 435) | 348 (287, 400) |
| | 30 | Thickness | 0.127 (0.093, 0.163) | 0.108 (0.0508, 0.162) | 0.144 (0.0994, 0.189) |
| | 30 | SLA | 20.6 (17, 24) | 22.3 (17.3, 27.2) | 20.5 (16.7, 24.1) |
| | 6 | Phosphorus | 2.31 (1.96, 2.67) | 2.33 (1.6, 3.04) | 1.96 (1.42, 2.72) |
